# Third Trimester-Equivalent Alcohol Exposure Induces Sex-Dependent Alterations in Locomotor Activity, Anxiety-Risky Behaviors, and Enhances Mechanical Allodynia in Adulthood

**DOI:** 10.64898/2026.04.28.721494

**Authors:** Estrella Villicana, Melody S. Sun, Haojie Chen, Luis E. Paez-Beltran, Carissa J. Milliken, Erik J. Balmer, Russell A. Morton, Erin Milligan, C. Fernando Valenzuela, Tou Yia Vue

**Affiliations:** Department of Neurosciences, University of New Mexico Health Sciences Center, Albuquerque, NM 87131; New Mexico Alcohol Research Center, University of New Mexico Health Sciences Center Albuquerque, NM 87131; Center for Brain Recovery & Repair Pre-Clinical Core, University of New Mexico Health Sciences Center Albuquerque, NM 87131

**Author notes:** **Corresponding Author**: Tou Yia Vue.

## Abstract

Prenatal alcohol exposure (PAE) causes fetal alcohol spectrum disorders (FASDs), which are neurodevelopmental conditions characterized by behavioral dysregulation, learning deficits, and cognitive inflexibilities. Alcohol exposure is harmful at all stages of human gestation, including the third trimester. This developmental window—characterized by rapid brain growth, myelination, and neural circuit formation—may be particularly vulnerable, yet the long-lasting behavioral and sensory consequences of exposure during this period remain poorly understood. In this study, neonatal mouse pups were exposed to ethanol (EtOH) or air vapor from postnatal day (P) 4 to P8, which is equivalent to a third-trimester alcohol exposure (TTAE) in humans. Blood ethanol concentrations measured at P8 reached approximately 250 mg/dL, consistent with binge-level exposure. Air- and EtOH-exposed mice were then assessed as adults at 5–6 months of age for locomotor activity, anxiety-related risky behaviors, recognition memory, and increased susceptibility to peripheral neuropathy, as indicated by sensitization to light touch following minor chronic constriction injury (mCCI) of the sciatic nerve. We found that TTAE was sufficient to produce long-lasting behavioral outcomes in a sex-dependent manner. Notably, EtOH-exposed males exhibited increased spontaneous locomotor activity and risky behavior, whereas EtOH-exposed females showed minimal or decreased changes compared to their respective controls. However, both EtOH-exposed male and female mice exhibited marked increases in light-touch sensitization, referred to as mechanical allodynia, following mCCI, a response absent in air-exposed controls. Together, these findings reveal that TTAE is highly detrimental to behavioral regulation and creates a vulnerability to developing neuropathic pain in adulthood.

## 1. INTRODUCTION

Prenatal alcohol exposure (PAE) remains a major cause of preventable neurodevelopmental injury that can lead to the development of fetal alcohol spectrum disorders (FASDs). FASD is characterized by a wide range of brain structural abnormalities, cognitive deficits, and behavioral inflexibilities due to the teratogenic effects of alcohol during pregnancy (Riley et al., 2011). Recent epidemiological studies estimate that FASDs affect approximately 1–5% of children in the United States, which is a substantial public health burden since individuals with FASDs frequently exhibit deficits in learning, executive function, emotional and sensory regulation, and adaptive behavior that extend into adolescence and adulthood (May et al., 2018; Mattson et al., 2019). Thus, FASDs impose significant societal and economic costs related to medical care, educational services, and requires long-term support and intervention (Glass et al., 2023), highlighting the need to identify critical windows of brain vulnerability and mechanisms underlying the neurodevelopmental and neurobehavioral consequences of these disorders .

In the last several decades, animal models have been essential for elucidating the developmental timing and cellular mechanisms through which alcohol disrupts brain maturation and function. In particular, rodent models have demonstrated that alcohol exposure during discrete developmental stages and severity of exposure produces distinct structural and behavioral outcomes, reflecting the stage-specific sensitivity of the developing brain to environmental insults (Goodlett et al., 2001; Guerri et al., 2009). The first and second trimesters corresponds to neurulation, proliferation of epithelial cells, neurogenesis and neuronal migration, while the third trimester represents a particularly critical period of increase in brain growth, volume, and maturation characterized by rapid synaptogenesis, myelination, and refinement of neural circuits required for sensory and behavioral development (Zhou et al., 2024). In rodents, prenatal development largely corresponds to the first and second trimesters of human gestation, whereas the human third trimester is developmentally equivalent to the early postnatal period, particularly the first two postnatal weeks, when rapid gliogenesis, synaptic maturation, myelination, and neural circuit refinement occur in the rodent brain (Semple et al, 2013). Although extensive work has characterized the consequences of alcohol exposure during early and mid-gestational development and considerable research has examined third-trimester-equivalent alcohol exposure (TTAE) (Guerri et al., 2009; Patten et al., 2014), important gaps remain in understanding the persistence and breadth of its effects later in life.

Given the vulnerability of glial development during late gestation, the present study investigated whether TTAE disrupts neurodevelopment in ways that produce enduring behavioral and somatosensory abnormalities. To address this, we employed a neonatal vapor exposure paradigm (Morton et al., 2014; Baculis et al., 2015) in which mouse pups were exposed to binge-level EtOH or air during the early postnatal period, coinciding with the peak of gliogenesis—a critical window when astrocyte and oligodendrocyte proliferation, migration, and differentiation are particularly susceptible to disruption (Guerri et al., 2001; Newville et al., 2017, 2022; Paez-Beltran et al., 2026). We then assessed adult outcomes across multiple domains relevant to FASD, including locomotor activity, exploratory and risk-related behaviors, cognitive performance, and mechanical sensitivity following minor peripheral nerve injury.

Our findings demonstrate that TTAE results in persistent behavioral alterations, characterized by increased exploratory and risk-taking behaviors and enhanced mechanical sensitivity in adulthood. Notably, these results provide new evidence that allodynia following late gestational alcohol exposure may be particularly pronounced. Collectively, these findings identify the third trimester as a vulnerable developmental window for long-lasting behavioral and sensory dysfunction and extend current understanding of how developmental alcohol exposure contributes to FASD-related phenotypes.

## 2. MATERIALS and METHODS

### 2.1. Animals

All mouse experiments in this study were conducted in compliance with the *NIH Guide for the Care and Use of Laboratory Animals* and were approved by the Institutional Animal Care and Use Committee (IACUC) at the University of New Mexico Health Sciences Center (UNM-HSC). Animals were housed in the UNM-HSC Animal Resource Facility (ARF). All mice were maintained on a 12-hour reverse light/dark cycle in a temperature-controlled environment during the mating, birth, and alcohol-exposure periods. Both control and experimental groups had 24-hour access to the same food (5V5R – PicoLab Select Rodent 50 IF/6F, LabDiet), water, and bedding.

### 2.2. Third Trimester-Equivalent Alcohol Exposure (TTAE) Paradigm

To model prenatal alcohol exposure (PAE) equivalent to the third trimester in humans, neonatal mice were exposed to either ethanol (EtOH) or air vapor from postnatal day (P) 4 to P8 using vapor chambers (Morton et al., 2014; Baculis et al., 2015) during the dark/awake phase (07:00 – 19:00). Each dam along with the pups were housed together in a standard cage with non-filter lid throughout the duration of the exposure to ensure adequate flow of vapors into the cage. On the first day of exposure, pups were exposed for 3 hours (10:00–13:00) to acclimate to the vapor chambers. For the remaining four days, mice were exposed for 4 hours daily (10:00–14:00). Breathalyzer measurements were obtained daily from each EtOH-exposed chamber to ensure consistent concentrations (7-8 g/dL). Blood ethanol concentration (BEC) was then collected from one pup per ethanol-exposed cage via rapid decapitation following the final exposure session to confirm appropriate levels of systemic alcohol. All pups and dams were then transferred to clean standard cages for maintenance.

### 2.3. Behavioral Studies – Cohort 1 Mice

All mice used for behavioral testing were between 5 and 6 months of age. Air- and EtOH-exposed experimental groups (N=8/group) consisted of approximately equal numbers of males and females from 4-5 different litters per group (see **Table 1**). Behavioral assessments were conducted in the following sequence across a period of about 30 days: Home Cage Activity Monitoring (Days 1-5), Open Field Test (Days 13), Novel Object Recognition (Days 16-17), and Elevated Zero Maze (Days 25). These tasks were spaced apart to ensure adequate recovery time while minimizing potential carryover effects and were performed between 12:00 and 16:00 during the light phase of the light-dark cycle similarly for both the male and female cohorts, which were run alternately on separate days.

**TABLE 1.**

| Cohort 1: Mice for Behavioral Studies |  |  |  |  |  |  |  |  | Cohort 2: Mice for Pain Studies |  |  |  |  |
| --- | --- | --- | --- | --- | --- | --- | --- | --- | --- | --- | --- | --- | --- |
| Litter ID | Mouse ID | Sex | Treatment | Home Cage | Open Field | Novel Object | EZM | CatWalk | Litter ID | Mouse ID | Sex | Treatment | Surgery |
| AL 30 | 10 | F | Air | ✓ | ✓ | ✓ | ✓ | ✓ | AL37 & 39 | 10 | F | Air | mCCI |
| AL 30 | 20 | F | Air |  | ✓ | ✓ | ✓ | ✓ | AL37 & 39 | 11 | F | Air | mCCI |
| AL 30 | 40 | F | Air | ✓ |  |  |  |  | AL41 | 31 | F | Air | mCCI |
| AL 30 | 60 | F | Air |  |  |  | ✓ |  | AL41 | 32 | F | Air | mCCI |
| AL 40 | 10 | F | Air |  | ✓ | ✓ | ✓ | ✓ | AL51 | 43 | F | Air | mCCI |
| AL 40 | 20 | F | Air | ✓ |  |  |  | ✓ | AL34 | 8 | F | EtOH | mCCI |
| AL 40 | 30 | F | Air |  |  | ✓ | ✓ |  | AL37 & 35 | 19 | F | EtOH | mCCI |
| AL 40 | 40 | F | Air |  | ✓ |  |  |  | AL37 & 35 | 22 | F | EtOH | mCCI |
| AL 44 | 10 | F | Air |  |  | ✓ |  | ✓ | AL37 & 35 | 23 | F | EtOH | mCCI |
| AL 44 | 20 | F | Air | ✓ | ✓ |  | ✓ |  | AL49 & 47 | 42 | F | EtOH | mCCI |
| AL 44 | 30 | F | Air | ✓ | ✓ | ✓ |  |  | AL37 & 39 | 9 | F | Air | Sham |
| AL 44 | 50 | F | Air |  |  |  | ✓ | ✓ | AL37 & 39 | 12 | F | Air | Sham |
| AL 45 | 10 | F | Air |  | ✓ |  | ✓ |  | AL41 | 33 | F | Air | Sham |
| AL 45 | 20 | F | Air |  | ✓ | ✓ |  | ✓ | AL51 | 44 | F | Air | Sham |
| AL 45 | 30 | F | Air |  |  |  |  | ✓ | AL51 | 45 | F | Air | Sham |
| AL 45 | 40 | F | Air | ✓ |  |  |  |  | AL34 | 7 | F | EtOH | Sham |
| AL 45 | 50 | F | Air |  |  | ✓ |  |  | AL37 & 35 | 20 | F | EtOH | Sham |
| AL 31 | 10 | F | EtOH | ✓ | ✓ | ✓ | ✓ | ✓ | AL37 & 35 | 21 | F | EtOH | Sham |
| AL 31 | 20 | F | EtOH | ✓ | ✓ | ✓ |  | ✓ | AL37 & 35 | 24 | F | EtOH | Sham |
| AL 33 | 10 | F | EtOH |  |  | ✓ | ✓ |  | AL49 & 47 | 41 | F | EtOH | Sham |
| AL 33 | 20 | F | EtOH | ✓ | ✓ |  | ✓ | ✓ | AL39 | 4 | M | Air | mCCI |
| AL 33 | 30 | F | EtOH |  | ✓ | ✓ | ✓ | ✓ | AL39 | 5 | M | Air | mCCI |
| AL 33 | 40 | F | EtOH | ✓ |  |  |  |  | AL36 | 25 | M | Air | mCCI |
| AL 32 | 10 | F | EtOH |  | ✓ |  | ✓ | ✓ | AL36 | 27 | M | Air | mCCI |
| AL 32 | 20 | F | EtOH | ✓ |  | ✓ | ✓ | ✓ | AL41 | 30 | M | Air | mCCI |
| AL 32 | 30 | F | EtOH | ✓ | ✓ | ✓ |  |  | AL51 | 49 | M | Air | mCCI |
| AL 38 | 10 | F | EtOH |  | ✓ | ✓ | ✓ | ✓ | AL34 | 1 | M | EtOH | mCCI |
| AL 38 | 20 | F | EtOH | ✓ |  | ✓ | ✓ |  | AL34 | 2 | M | EtOH | mCCI |
| AL 38 | 30 | F | EtOH |  | ✓ |  |  | ✓ | AL37 & 35 | 15 | M | EtOH | mCCI |
| AL 30 | 10 | M | Air | ✓ | ✓ |  | ✓ |  | AL37 & 35 | 16 | M | EtOH | mCCI |
| AL 30 | 20 | M | Air |  | ✓ |  |  | ✓ | AL37 & 35 | 17 | M | EtOH | mCCI |
| AL 30 | 30 | M | Air |  |  | ✓ | ✓ |  | AL43 | 34 | M | EtOH | mCCI |
| AL 30 | 40 | M | Air | ✓ |  | ✓ |  | ✓ | AL43 | 35 | M | EtOH | mCCI |
| AL 40 | 10 | M | Air | ✓ | ✓ | ✓ | ✓ | ✓ | AL43 | 37 | M | EtOH | mCCI |
| AL 40 | 20 | M | Air |  | ✓ | ✓ | ✓ | ✓ | AL47 | 40 | M | EtOH | mCCI |
| AL 42 | 10 | M | Air | ✓ | ✓ | ✓ | ✓ | ✓ | AL39 | 6 | M | Air | Sham |
| AL 42 | 20 | M | Air |  | ✓ | ✓ | ✓ | ✓ | AL36 | 26 | M | Air | Sham |
| AL 45 | 10 | M | Air |  | ✓ | ✓ | ✓ | ✓ | AL36 | 28 | M | Air | Sham |
| AL 45 | 20 | M | Air | ✓ | ✓ |  |  | ✓ | AL41 | 29 | M | Air | Sham |
| AL 45 | 30 | M | Air | ✓ |  | ✓ | ✓ |  | AL51 | 48 | M | Air | Sham |
| AL 31 | 10 | M | EtOH | ✓ | ✓ | ✓ |  | ✓ | AL34 | 3 | M | EtOH | Sham |
| AL 31 | 20 | M | EtOH | ✓ |  | ✓ | ✓ | ✓ | AL37 & 35 | 13 | M | EtOH | Sham |
| AL 33 | 10 | M | EtOH |  | ✓ | ✓ | ✓ | ✓ | AL37 & 35 | 14 | M | EtOH | Sham |
| AL 32 | 10 | M | EtOH | ✓ | ✓ |  | ✓ | ✓ | AL37 & 35 | 18 | M | EtOH | Sham |
| AL 32 | 20 | M | EtOH | ✓ |  | ✓ |  |  | AL43 | 36 | M | EtOH | Sham |
| AL 38 | 10 | M | EtOH | ✓ | ✓ | ✓ | ✓ | ✓ | AL43 | 38 | M | EtOH | Sham |
| AL 38 | 20 | M | EtOH |  |  | ✓ |  |  | AL47 | 39 | M | EtOH | Sham |
| AL 38 | 30 | M | EtOH |  | ✓ |  | ✓ | ✓ | AL49 | 46 | M | EtOH | Sham |
| AL 46 | 10 | M | EtOH |  | ✓ | ✓ | ✓ |  | AL49 | 47 | M | EtOH | Sham |
| AL 46 | 20 | M | EtOH | ✓ | ✓ |  | ✓ | ✓ |  |  |  |  |  |
| AL 46 | 30 | M | EtOH | ✓ | ✓ | ✓ | ✓ | ✓ |  |  |  |  |  |

#### 2.3.1. Home Cage Activity Monitoring – Days 1-5

Mice were single-housed in standard cages with corncob bedding and monitored for five consecutive days (112 hours) to establish baseline locomotor activity. Food and water were available ad libitum. The first day/24 hour served as an acclimation period to the new environment. Each cage was placed within a metal-framed enclosure equipped with photoelectric beam sensors (PAS Home Cage System; San Diego Instruments) that recorded locomotor activity based on beam breaks. Activity data were collected continuously across both light (inactive) and dark (active) phases. Cages were checked daily to prevent sensor obstruction, but those with sensors blocked by bedding or that malfunctioned for a significant period were excluded from the analysis.

#### 2.3.2. Open Field Test – Day 13

The Open Field test was used to assess anxiety-like behavior and exploratory activity. At the start of the task, each mouse was placed in the center of a white Plexiglas arena and allowed to explore freely for 10 minutes. A ceiling-mounted camera (Med Associates Basler acA1300–60) and EthoVision XT tracking software (Noldus Information Technology, Netherlands) were used to monitor movement and quantify time spent in the center versus border zones. To minimize olfactory cues, the arena was cleaned with 70% ethanol between trials.

#### 2.3.3. Novel Object Recognition Task – Days 16-17

The NOR task was used to assess memory and anxiety-like behavior. The task was conducted in the same arena used for the OFT. On day one (familiarization phase), each mouse was placed in the center of the arena between two identical objects and allowed to explore for 10 minutes. On day two (test phase), the mouse was reintroduced to the arena for 10 minutes and presented with one familiar object and one novel object.

Mouse interactions with the objects were recorded using an overhead camera and analyzed with EthoVision XT software. Both the arena and objects were cleaned with 70% ethanol between trials to eliminate olfactory cues.

#### 2.3.4. Elevated Zero Maze Task – Day 25

The EZM was used to assess anxiety-like and risky behavior. The apparatus consisted of an elevated circular track divided into two closed (enclosed) arms and two open (exposed) arms. The closed arms provided shelter from light and external stimuli, whereas the open arms exposed mice to bright light and the surrounding environment.

Each mouse was initially placed on the same open arm and allowed to explore freely for 10 minutes. Movement and arm preference were recorded using an overhead camera and analyzed with EthoVision XT software.

#### 2.4. Pain Studies – Cohort 2 Mice

All air- and EtOH-exposed mice used were between 5-6 months of age and consisted of both females and males from 3-4 different litters per group (see Table 1).

#### 2.4.1. Minor Chronic Constriction Injury (mCCI) Surgery

Mechanical allodynia, which is a peripheral neuropathic condition, was induced using a minor chronic constriction injury (mCCI) of the sciatic nerve previously shown to induce mechanical allodynia only in offspring with PAE (Sanchez et al., 2017; Noor et al., 2019). The mCCI model induces a mild peripheral nerve injury through loose ligation of one sciatic nerve, enabling reliable assessment of hind paw threshold responses to light touch with minimal impact on motor function. This approach is particularly well-suited for detecting mechanical allodynia, defined as pain in response to a normally non-noxious stimulus.

All procedures were performed as previously detailed, with minor modifications (Noor et al., 2019). Briefly, mice were anesthetized with isoflurane (1.5–2.0% in oxygen; induction at 5.0 L/min, surgical plane at 2.0 L/min) and placed on a sterile surgical field. The dorsal surface of the left thigh was shaved and disinfected with 70% ethanol prior to incision. Using aseptic technique, a small incision (∼2 cm) was made in the skin just caudal to the femur using a sterile scalpel blade, followed by blunt dissection through the muscle fascia to expose the sciatic nerve. The sciatic nerve was carefully isolated from surrounding tissue using sterile 200 ul pipette tips as probes. A single ligature of 6-0 chromic gut suture (Ethicon: SUTURE, 6/0 18 CHROMIC GUT, BL G-6, VA) was gently tied around the sciatic nerve proximal to the trifurcation. The ligature was secured to produce a slight constriction of the nerve without overt compression or interruption of epineural blood flow. Throughout the procedure, the exposed nerve and surrounding tissues were intermittently irrigated with sterile isotonic saline to prevent dehydration. Following ligation, the sciatic nerve was returned to its anatomical position, and the muscle fascia was closed using a single 4-0 silk suture. The skin incision was closed using sterile wound clips, which were removed 7 days after surgery. Sham-operated mice underwent identical surgical procedures, including nerve exposure, but without chromic gut suture ligation. All surgical procedures were completed within approximately 15–20 minutes. Mice recovered from anesthesia within approximately 10 minutes and were returned to their home cages. Animals were monitored daily for postoperative recovery and general health status, including body weight, wound condition, hind paw integrity, locomotor activity, and grooming behavior. All mice recovered and remained in the study. The number of mice.

#### 2.4.2. Assessment of Mechanical Allodynia (von Frey Test)

Mechanical allodynia was assessed using calibrated von Frey monofilaments applied to the plantar surface of the hind paw, as previously described for mouse neuropathic pain models (Vanderwall et al., 2018; Noor et al., 2019). Behavioral testing was conducted during the first 3 hours of the light phase of the light–dark cycle.

Prior to baseline testing, mice were habituated to the experimenters and testing environment for 45–60 minutes per day for four consecutive days. Baseline mechanical sensitivity was then measured once per day, with assessments separated by at least 48 hours. For testing, mice were placed individually in transparent chambers positioned on an elevated mesh platform and allowed to acclimate before stimulation. Mechanical stimuli were delivered using nine calibrated von Frey monofilaments applied perpendicularly to the plantar surface of the hind paw for a maximum duration of 3 seconds. Inter-stimulus intervals were maintained at approximately 30 seconds to minimize sensitization.

The set of monofilaments spanned the following force range: the log intensity of the nine monofilaments used is defined as log10 (grams × 10,000) with the range of intensity being as follows, reported in log (grams): 2.36 (0.022 g), 2.44 (0.028 g), 2.83 (0.068 g), 3.22 (0.166 g), 3.61 (0.407 g), 3.84 (0.692 g), 4.08 (1.202 g), 4.17 (1.479 g), and 4.31 (2.042 g). Testing began with the 3.22 log unit filament (0.166 g), corresponding to the midpoint of the stimulus range. Subsequent filament selection followed an up–down method based on the mouse’s response to the previous stimulus. If no withdrawal response occurred, the next higher-force filament was applied. If a withdrawal response was observed, the next lower-force filament was used. This sequence continued for up to six stimulus applications per paw.

Withdrawal responses included rapid paw withdrawal, shaking, or licking. Both the ipsilateral (injured) and contralateral (uninjured) hind paws were tested during each session, with testing order randomized to minimize order effects. Mechanical sensitivity thresholds were recorded longitudinally following surgery at predetermined time points selected based on prior studies demonstrating the onset and progression of neuropathic hypersensitivity in the mCCI model.

### 2.4. Statistical Analysis

Behavioral data from Cohort 1 mice were analyzed using statistical tests appropriate to the experimental design of each behavioral assay. For comparisons involving a single independent factor, unpaired t-tests with Welch’s correction were used. For assays involving two independent variables, data were analyzed using 2-way analyses of variance (ANOVA) with Sex and Treatment as between-subject factors. When applicable, interactions between factors (e.g., Sex × Treatment) were evaluated. Significant main effects or interactions were followed by uncorrected Fisher’s least significant difference (LSD) post hoc multiple comparisons tests to compare individual group means.

For Cohort 2 mice, mechanical sensitivity data obtained from the von Frey test following mCCI were analyzed using two-way repeated-measures (RM) ANOVA, with Time (baseline and days post-injury) as the within-subject factor and Treatment (Air vs. EtOH) as the between-subject factor. Individual animals were treated as repeated measures (subjects) across time. When the assumption of sphericity was violated, the Geisser–Greenhouse correction was applied to adjust degrees of freedom. Significant main effects or interactions were followed by Šídák’s multiple comparisons tests to compare treatment groups (i.e., Air mCCI vs EtOH mCCI) at individual time points.

All data are presented as mean ± SEM, and statistical significance was defined as p < 0.05. Statistical analyses were performed using GraphPad Prism (version 11.0; GraphPad Software, San Diego, CA, USA).

## 3. RESULTS

### 3.1. Third-trimester-equivalent alcohol exposure (TTAE) using vapor chamber produces binge-level ethanol exposure

The third trimester in humans corresponds to a period of neurite outgrowth, extensive gliogenesis, followed by synaptogenesis and the onset of myelination of major axon tracts, resulting in a rapid increase in the volume of the developing fetal brain (Zhou et al., 2024). In rodents, this third trimester-equivalent developmental period occurs during the first two postnatal weeks prior to eye opening. To determine the long-term consequences of alcohol exposure during the third trimester on behavioral and sensory modalities, we exposed neonatal mice to either air or EtOH vapor for four hours (10:00–14:00) per day under a reverse light cycle from postnatal day (P) 4 to P8 using a vapor chamber system (**Fig. 1a**) (Morton et al., 2014; Baculis et al., 2015). This exposure paradigm was selected because it produces consistent EtOH exposure across multiple litters simultaneously while minimizing direct handling of mouse pups and dams throughout the exposure period. Blood ethanol concentration (BEC) was assessed using one pup per EtOH-exposed litter on the final day of exposure (P8) and demonstrated robust systemic levels averaging approximately 250 mg/dL (**Fig. 1b**), consistent with binge-level alcohol exposure.

**Figure 1.**
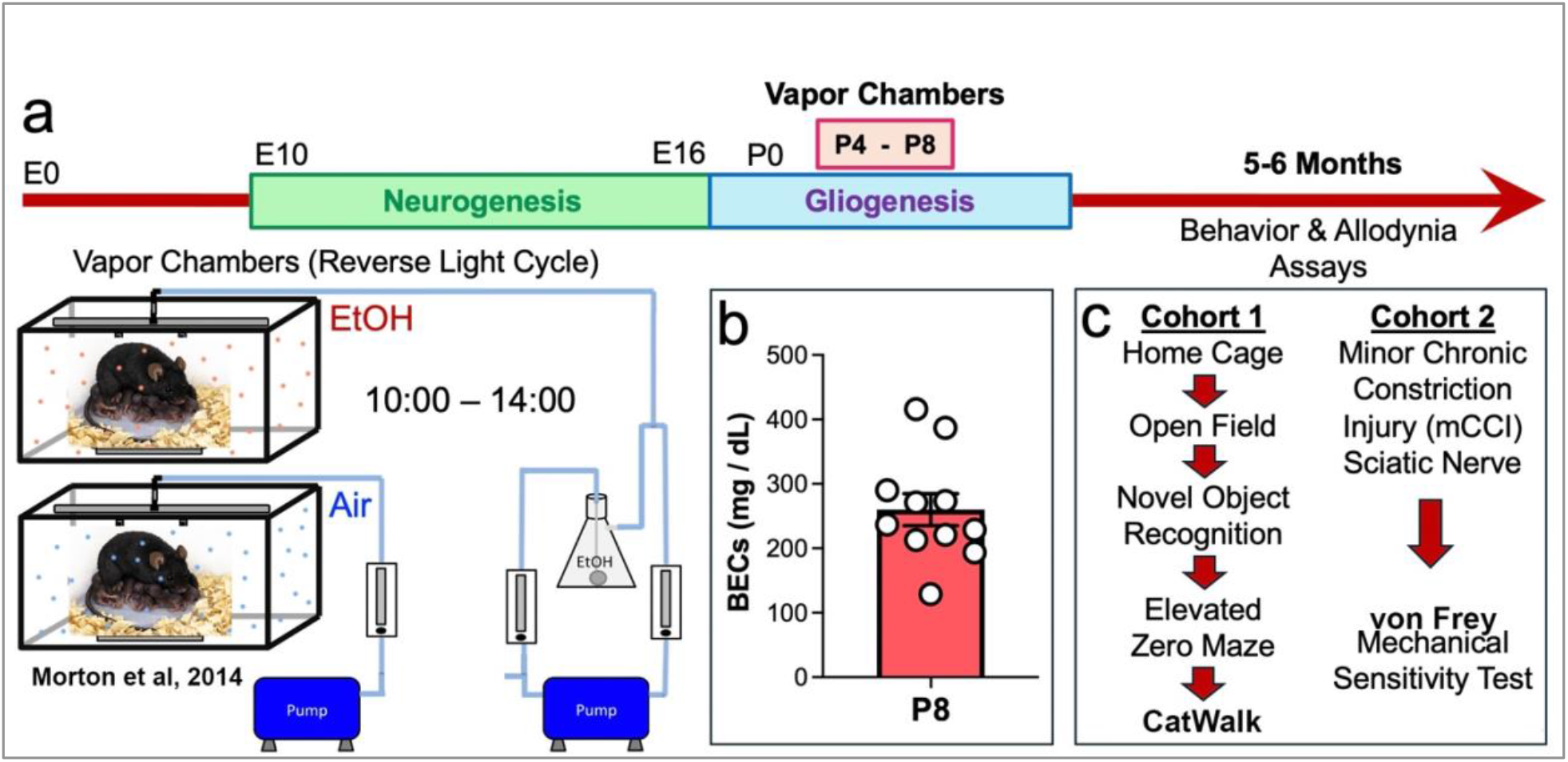
Third-trimester-equivalent alcohol exposure (TTAE) paradigm and behavioral assays. **(a)** Experimental timeline illustrating vapor chamber exposure TTAE paradigm. Females with neonatal mouse pups were exposed to EtOH or Air vapor from postnatal day (P)4 to P8 for 4 hours per day under a reverse light cycle (10:00–14:00). This developmental window corresponds to late gestation in humans and represents a critical period of brain growth and maturation. Following completion of the exposure paradigm, animals were allowed to mature to adulthood for behavioral testing at 5-6 months of age. **(b)** Blood ethanol concentrations (BECs) were measured at P8 immediately following the final exposure session. Each data point represents 1 pup from each EtOH-exposed litter. Data are presented as mean ± SEM. **(c)** Two independent cohorts of male and female mice exposed to EtOH or Air vapor were used in this study. Cohort 1 was used for the behavior assays indicated to evaluate baseline locomotor activity, recognition-memory, anxiety-related risky behaviors, and gait & locomotion. Cohort 2 was used to assess susceptibility to neuropathic pain with minor chronic constriction injury (mCCI) of the sciatic nerve followed by mechanical sensitivity testing using von Frey filaments.

To assess long-term behavioral consequences, both EtOH-exposed and air-exposed control groups were tested in adulthood between 5–6 months of age. Two independent cohorts consisting of both males and females were subjected to a series of behavioral and sensory assays (**Fig. 1c**). Cohort 1 was used to assess baseline locomotor activity, exploratory and recognition memory, and anxiety- and risk-related behaviors sequentially across a one-month period beginning with Home Cage monitoring (days 1–5), followed by the Open Field Test (day 13), Novel Object Recognition (days 16–17), and the Elevated Zero Maze (day 25). Cohort 2 was used to determine susceptibility to neuropathic pain following peripheral nerve injury using the minor chronic constriction injury (mCCI) model and von Frey mechanical sensitivity testing (**Table 1**). All experiments within each cohort were performed by the same experimenter.

### 3.2. TTAE increases locomotor activity in a sex-dependent manner while preserving circadian rhythmicity

To assess spontaneous locomotor behavior, EtOH- or Air-exposed mice were single-housed in standard cages containing corncob bedding and placed within a metal-framed enclosure equipped with multiple photoelectric beam sensors arranged parallel and perpendicular to the cage. In this monitoring system, movement of the mice generated beam breaks that were recorded continuously for 5 days (∼112 hours total) across multiple light (sleep: 19:00–07:00) and dark (awake: 07:00–19:00) phases. The first day (24 hours) served as an acclimation period to the new cage environment, after which beam breaks were recorded and analyzed for all mice during days 2-4 (24-96 hours) (**Fig. 2a**).

**Figure 2.**
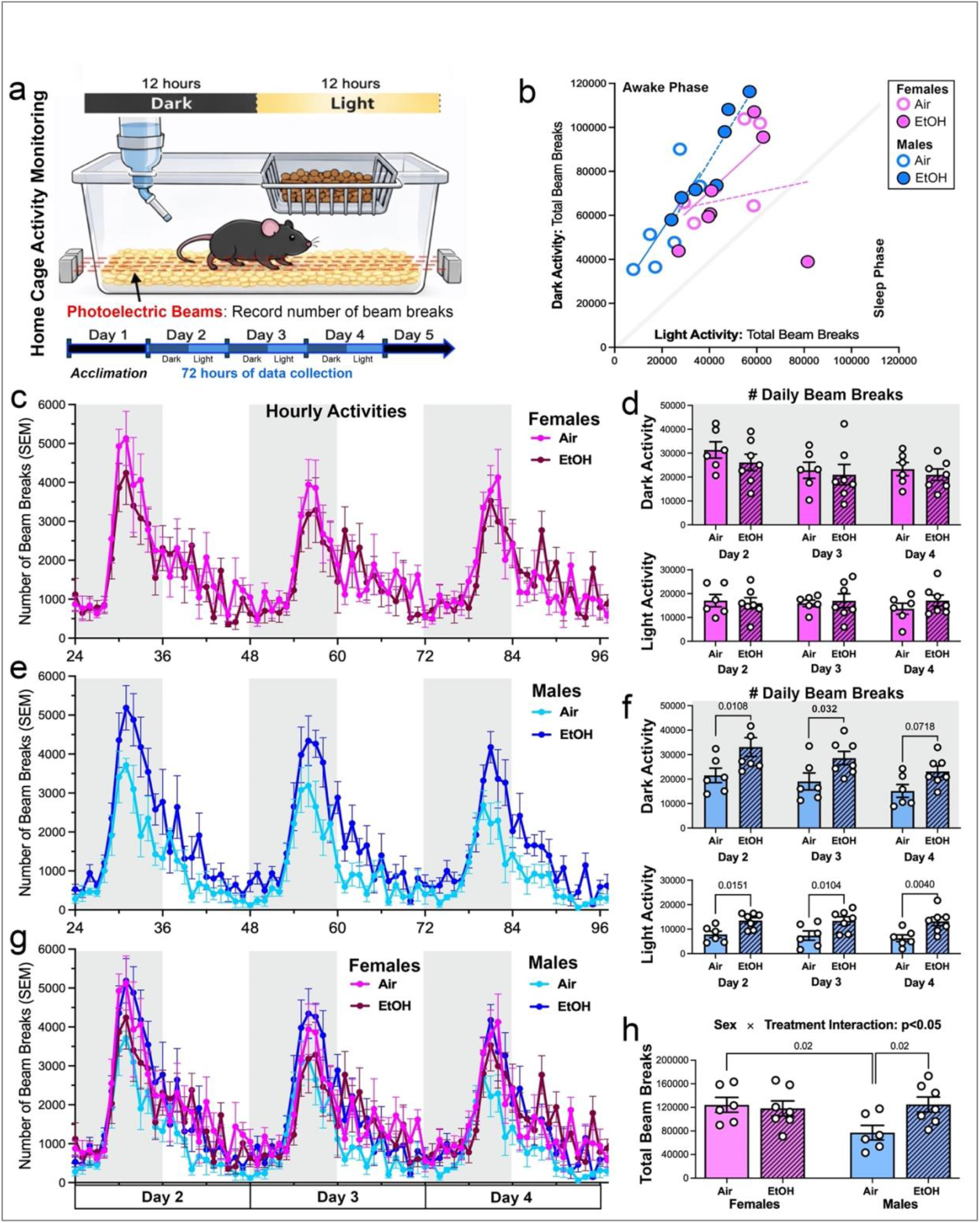
Developmental alcohol exposure increases spontaneous locomotor activity in a sex-dependent manner. **(a)** Schematic of a home cage monitoring system using photoelectric beam detection. Mice were singly housed and allowed to acclimate for 24 hours, followed by continuous recording of locomotor activity (beam breaks) for 72 hours across alternating 12-hour dark (19:00 – 07:00) and 12-hour light (07:00 – 19:00) cycles. **(b)** Scatter correlation plot comparing total beam breaks during dark (awake/active) and light (sleep/inactive) phases across male and female groups. All mice exhibited greater activity/total beam breaks during the dark/active phase except for one EtOH-exposed female. Solid correlation slope lines are for air-exposed groups while dashed correlation slopes are for EtOH-exposed groups. (**c, e, g**) Hourly beam breaks across Days 2-4 (72 hours) for females (c), males (e), and combined groups (g). Gray and light shading indicate dark and light phases, respectively. Data are presented as mean ± SEM. (**d, f**) Quantification of daily beam breaks during dark and light phases. Data are presented as mean ± SEM with individual data points. Statistical significance (p-values shown) was determined using 2-way ANOVA with multiple comparisons. Males (but not females) showed significance of main effect of treatment during both dark and light cycles. Dark activity: Treatment, F(1, 33) = 15, p = 0.0004. Light activity: Treatment, F(1, 33) = 23, p < 0.0001. (**h**) Quantification of total beam breaks for Days 2-4. Data are presented as mean ± SEM. Sex × treatment interactions (F(1, 22) = 4.4, p = 0.459) were determined using 2-way ANOVA. The effect of sex (air-exposed female vs male control groups) and the effect of treatment (Air-exposed vs. EtOH-exposed males) were determined using *t*-test with Welch’s correction.

A scatter plot analysis of total beam breaks for individual mice was first performed to determine whether any abnormal activity patterns occurred during the light and dark phases across the three recording days. This analysis revealed that nearly all mice behaved as expected for nocturnal animals, with total beam breaks consistently higher during the dark/awake phase than during the light/sleep phase. Positive correlation slopes illustrated that mice that were generally more active during the dark/awake cycle were also more active during the light/sleep cycle. However, one EtOH-exposed female displayed an atypical reverse activity pattern, causing approximately twice as many beam breaks during the light/sleep phase compared to the dark/awake phase (**Fig. 2b**).

We next analyzed hourly beam break patterns, which demonstrated that both male and female groups, regardless of treatment, exhibited clear circadian rhythms in locomotor activity, with peak beam breaks occurring during the midpoint of the dark phase followed by a progressive decline into the light phase (**Fig. 2c-e**). Despite this intact circadian rhythmicity, EtOH exposure induced sex-dependent alterations in locomotor activity. Specifically, EtOH-exposed females (dark red line) displayed hourly beam break patterns that closely resembled those of Air-exposed females (pink line) across all three recording days, although with modest reductions in peak activity during the dark phase (**Fig. 2c**). Despite this slight reduction, daily beam break counts revealed no significant differences for effect of treatment or days between EtOH- and air-exposed females across either dark or light phases during the three recording days (**Fig. 2d**), indicating that TTAE has minimal impact on female locomotor activity.

In contrast, EtOH-exposed males (dark blue line) exhibited consistently elevated hourly beam breaks compared to Air-exposed males (light blue line) throughout the three-day recording period (**Fig. 2e**). Daily beam break counts revealed that there was significant effect of treatment during both dark (F(1, 33) = 15, p=0.0004) and light (F(1, 33) = 23, p<0.0001) phases. Specifically, direct comparison on a day-by-day basis showed that there was significant increase in daily activity for EtOH-exposed males during the dark phase on Days 2 and 3 and across all three days during the light phase (**Fig. 2f**). Notably, locomotor activity levels in EtOH-exposed males reached values comparable to those observed in female groups, both of which were higher than those of air-exposed control males (**Fig. 2g, h**). Furthermore, 2-way ANOVA analysis of total beam breaks across all three recorded days revealed a significant Sex × Treatment interaction (F(1, 22) = 4.4, p<0.05) (**Fig. 2h**), indicating that TTAE produces persistent hyperactivity-like behavior selectively in males while exerting minimal effects on locomotor activity in females.

### 3.3. TTAE does not alter baseline anxiety-like behavior in the Open Field Test

To determine whether increased locomotor activity reflected changes in anxiety-like or exploratory behavior, animals were tested in the open field arena (**Fig. 3a**). Mice were placed in the center of the arena and allowed to explore freely for 10 minutes (600 seconds). Heatmap analyses revealed that animals from all groups displayed typical exploratory patterns characterized by a predominant preference for the corners and perimeter walls, with intermittent entries into the center zone (**Fig. 3b**).

**Figure 3.**
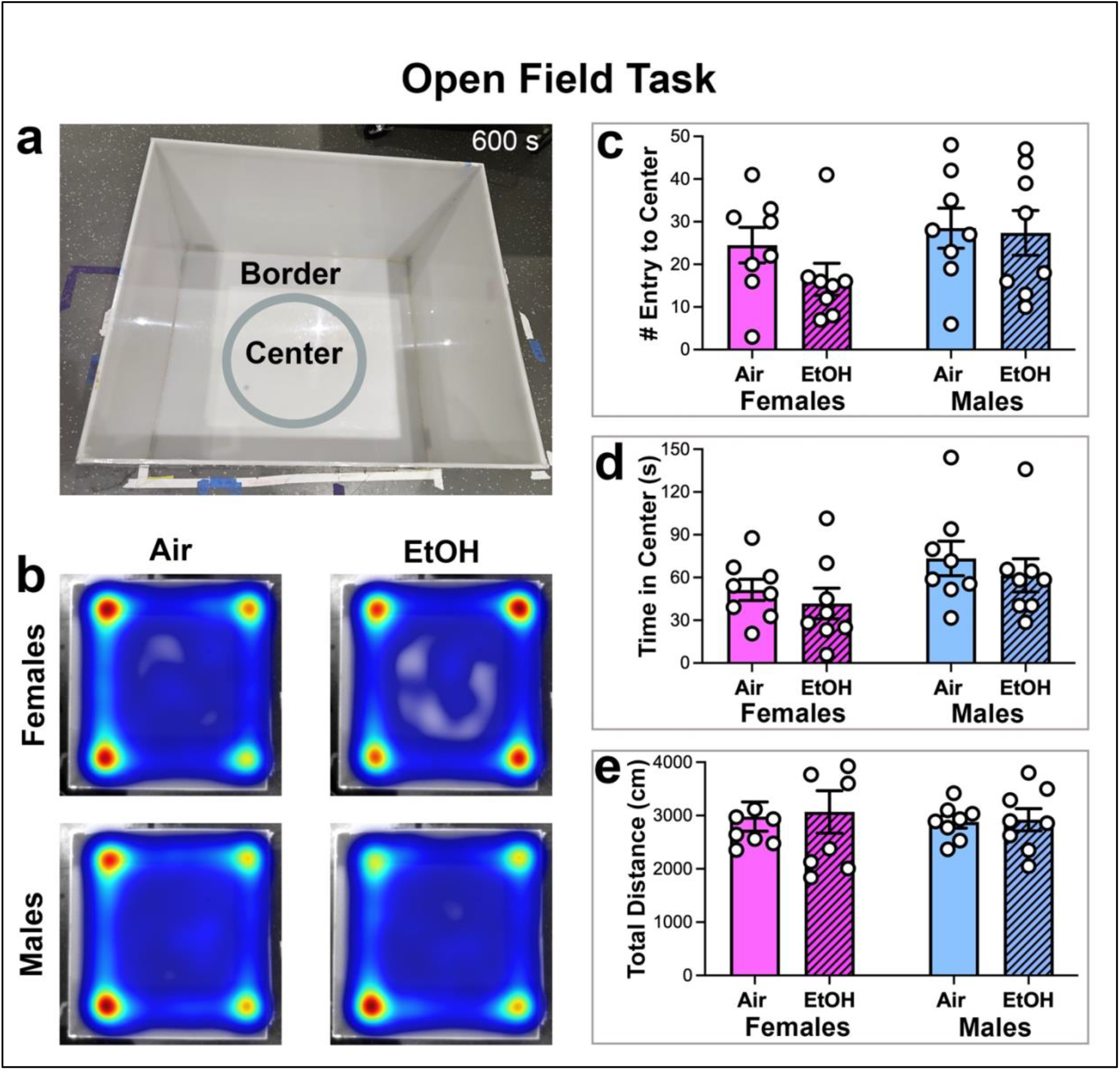
TTAE does not alter exploratory behavior in the Open Field test. **(a)** Open field arena with defined center and border zones in which mice were placed in the center zone and allowed to explore for 10 minutes. **(b)** Average heatmaps illustrating spatial exploration patterns across treatment groups. Note that males show greater exploration of center zone. (**c-e**) Number of entry (c) and time spent (d) in the center zone and total distance traveled (e). Data are presented as mean ± SEM with individual data points shown. Statistical analysis of effect of sex, treatment, or Sex × Treatment interactions using 2-way ANOVA with multiple comparisons showed no significance.

Quantification of center exploration frequency revealed no significant differences between air- and EtOH-exposed groups for either sex (**Fig. 3c**). Similarly, total time spent in the center zone during the 600-second testing period was comparable between treatment groups. However, on average, male mice were more likely to enter and remain in the center than female mice, with EtOH-exposed females showing the lowest willingness to explore or remain in the center zone (**Fig. 3d**). Despite these behavioral tendencies, all groups traveled comparable total distances within the arena (**Fig. 3e**).

Taken together, these findings indicate that TTAE does not produce overt baseline anxiety-like behavior in adulthood and suggest that the increased locomotor activity observed in male mice reflects altered locomotor regulation rather than generalized anxiety.

### 3.4. TTAE impairs recognition memory primarily in male mice

We next evaluated recognition memory using the novel object recognition task conducted within the same Open Field arena. During the training session on Day 1, mice were individually placed in the arena containing two identical objects positioned in opposite corners and allowed to explore for 10 minutes (**Fig. 4a**). After a 24-hour retention interval (Day 2), mice underwent the same exploration paradigm, except that one familiar object was replaced with a novel object to assess recognition memory (**Fig. 4b**).

**Figure 4.**
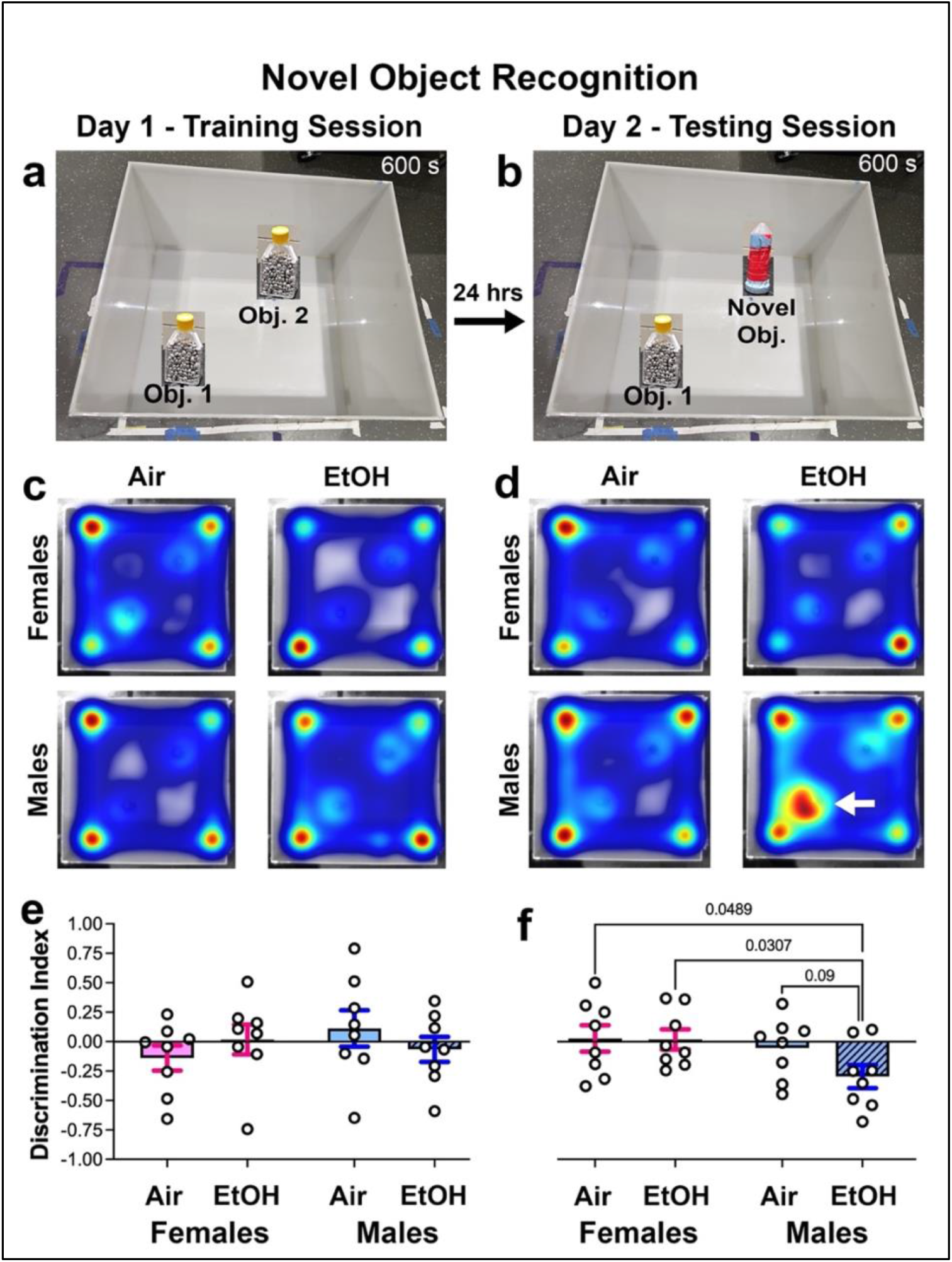
TTAE alters recognition memory in male mice. **(a)** Arena with two identical objects used for the training session during Day 1. **(b)** Same arena for the testing session 24 hours later with Object 2 replaced by a Novel Object. (**c, d**) Heatmaps of exploration patterns and time spent with objects on Day 1 (c) and Day 2 (d). Arrow indicate EtOH males showed increased interaction with familiar/Object 1 on Day 2. (**e, f**) Discrimination index for Day 1 (e) and Day 2 (f). DI = (Time w/ Obj. 2/N.O – Time w/ Obj. 1)/(Total Time w/ Obj. 1+2/N.O). Data are presented as mean ± SEM. Statistical analysis was performed using unpaired *t*-test with Welch’s correction.

Heatmap analyses revealed similar exploration patterns during the Day 1 training session across all treatment groups in both sexes, indicating equivalent interaction with the identical objects (**Fig. 4c**). During the Day 2 testing session, both EtOH- and air-exposed females, as well as air-exposed males, exhibited comparable interaction times with the familiar and novel objects. In contrast, EtOH-exposed males unexpectedly spent more time interacting with the familiar object than with the novel object (**Fig. 4d**).

Quantitative analysis confirmed these observations using the discrimination index (DI: time with novel object ™ time with familiar object / total time with both objects) (Antunes et al., 1998). Specifically, discrimination index values during the Day 1 training session were near zero across all groups, reflecting equivalent interaction with both identical objects (**Fig. 4e**). Following the 24-hour retention interval on Day 2, discrimination index values remained near zero for both female groups and air-exposed males, indicating no preference for the novel object. By contrast, EtOH-exposed males exhibited a negative discrimination index (∼™0.3), indicating a clear preference for the familiar object over the novel one (**Fig. 4f**).

These findings, particularly the absence of increased interaction with the novel object, were unexpected and may partially reflect the relatively long 24-hour retention interval between training and testing sessions, which may have reduced memory retention across groups. Nonetheless, the observation that recognition memory deficits were selectively evident in EtOH-exposed males but not females further highlights the persistent sex-specific behavioral consequences of TTAE.

### 3.5. TTAE produces sex-specific alterations in risk-related behavior in the elevated zero maze

To further assess anxiety-like behavior and risk evaluation under environmental exposure conditions, mice were tested in the elevated zero maze (**Fig. 5a**). Animals were placed in one of the open arms and allowed to explore the apparatus freely for 10 minutes. Heatmap analyses demonstrated that mice from all groups actively explored both open and closed segments of the maze (**Fig. 5b**), with greater time spent in the closed arms, indicating intact locomotor function and preserved exploratory behaviors across experimental groups.

**Figure 5.**
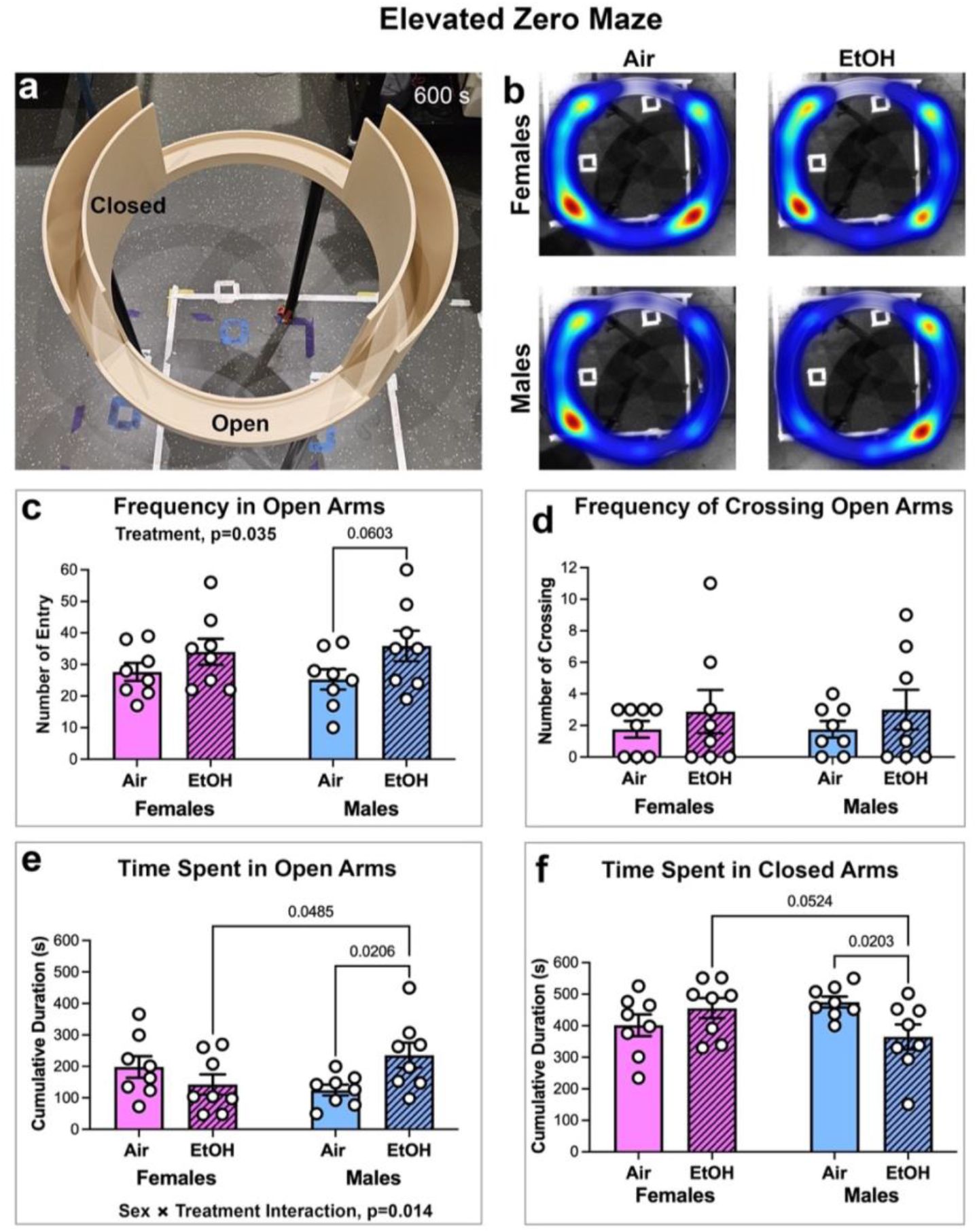
TTAE increased risky behavior in male mice. **(a)** Elevated zero maze apparatus consisting of alternating open and closed arms. Mice were placed in open arms and allowed to explore for 10 minutes. **(b)** Averaged heatmaps illustrating spatial exploration patterns across treatment groups. (**c, d**) Frequency of entries into (c) and crossing of open arms. (**e, f**) Time spent (d) in open arms or closed arm border zones during the 600-second testing period. Data are presented as mean ± SEM with individual data points shown. Statistical analysis of the effect of Treatment (F(1, 28) = 4.9, p = 0.035) and Sex x Treatment interaction (F(1, 28) = 6.8, p = 0.014) was determined using 2-way ANOVA with multiple comparisons.

Quantitative analysis revealed a significant effect of treatment (F(1, 28) = 4.9, p = 0.035) on the frequency of entries into the open arms, with EtOH-exposed mice exhibiting a greater number of entries into the open arms compared to air-exposed controls (**Fig. 5c**). Consistent with this pattern, a subset of EtOH-exposed mice displayed high-frequency traversal behavior, with approximately 30% (5/16) completing five or more full crossings of the open arms, whereas none of the Air-exposed controls exceeded four full crossings (**Fig. 5d**). This elevated exploratory drive suggests that TTAE reduces behavioral inhibition into open, high-risk environment in a subset of animals.

In contrast, 2-way ANOVA analysis of cumulative time spent in the open arms revealed a significant Sex × Treatment interaction (F(1, 28) = 6.8, p = 0.014), indicating that the behavioral consequences of TTAE differed between males and females (**Fig. 5e**). Specifically, EtOH-exposed females spent less time in the open arms (approximately 140 s; ∼25% of total time) compared to air-exposed female controls (approximately 200 s; ∼33%), suggesting a modest increase in avoidance of exposed, high-risk environments. In contrast, EtOH-exposed males spent the greatest amount of time in the open arms (approximately 235 s; ∼40%), representing a two-fold increase relative to air-exposed control males (approximately 125 s; ∼20%; p = 0.0206). A direct comparison between EtOH-exposed groups further confirmed that males also spent significantly more time (p = 0.0485) in the open arms than the EtOH-exposed females (**Fig. 5e**). Conversely, the EtOH-exposed females spent significantly more time within the closed arms than the EtOH-males (p=0.05) (**Fig. 5f**).

Taken together, these findings demonstrate that TTAE produces a sex-dependent divergence in risk-related behavior. Specifically, while TTAE appears to increase overall exploratory drive, risk-taking behavior is enhanced in males but reduced in females.

### 3.6. TTAE increases susceptibility to neuropathic pain following peripheral nerve injury

Rodent models of prenatal alcohol exposure during the first and second trimesters have been shown to develop adult neuropathic pain when challenged with minor injury (Noor et al., 2020). Whether a similar vulnerability exists following third-trimester-equivalent exposure remains unknown. To address this question, a second cohort of air- or EtOH-exposed mice underwent mCCI of the sciatic nerve, followed by assessment of mechanical sensitivity using von Frey filaments (**Fig. 6a,b**). This model produces chronic neuropathy that enables the detection of mechanical allodynia, which is absent in sham surgery controls and non-exposed animals.

**Figure 7.**
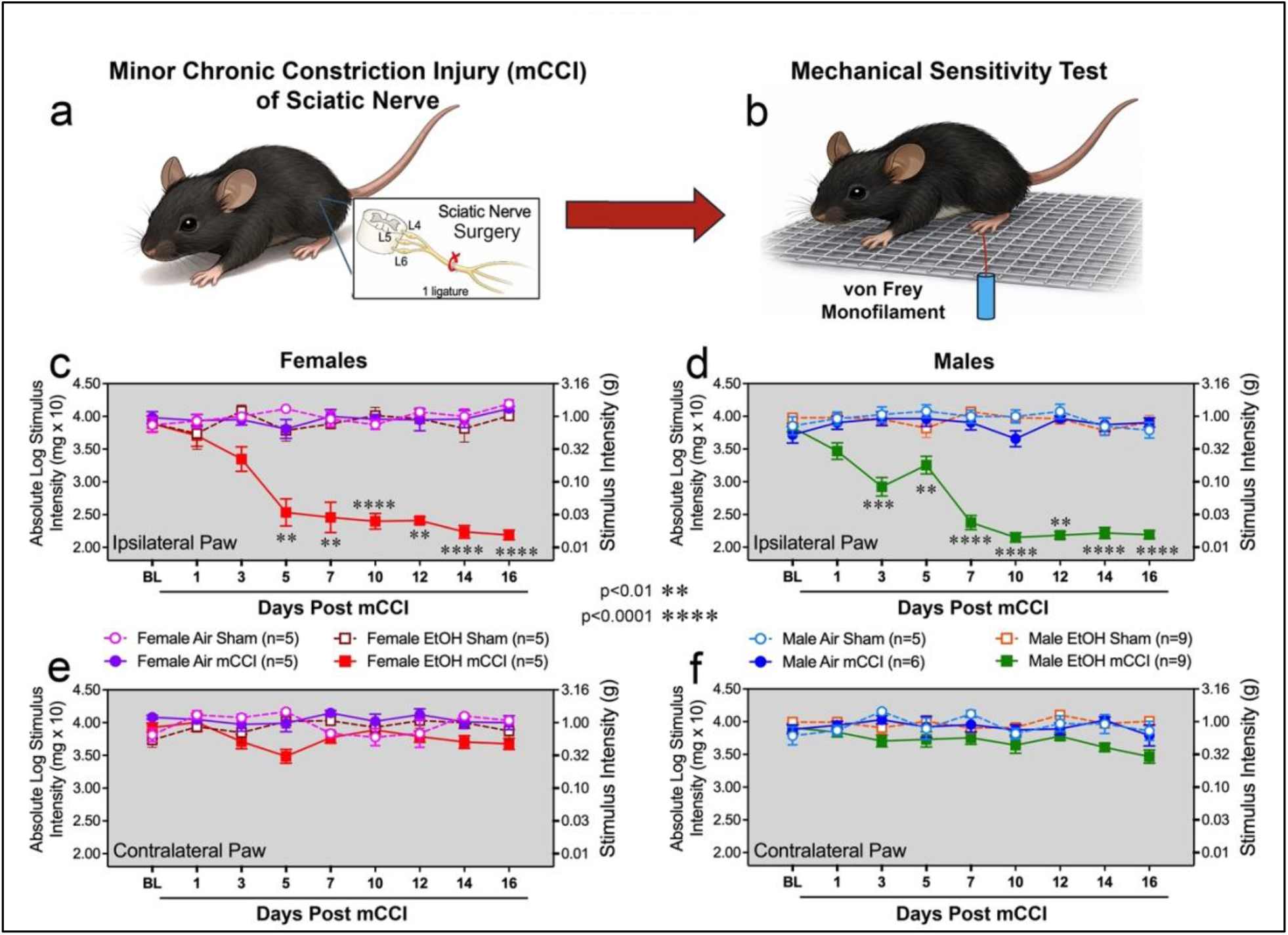
Developmental alcohol exposure increases susceptibility to neuropathic pain following minor chronic constriction injury. (**a, b**) Experimental schematic illustrating minor chronic constriction injury (mCCI) of the sciatic nerve (a) followed by mechanical sensitivity testing using von Frey filaments (b). Sham control mice underwent a similar surgical procedure but were not subjected to the single ligature constriction. (**c, d**) Mechanical withdrawal thresholds measured from the ipsilateral paw following injury in female (c) and male (d) mice showing allodynia in EtOH-exposed mCCI female and male mice but not Air-exposed or Sham control mice. (**e, f**) Mechanical withdrawal thresholds measured from the uninjured contralateral paw of female (e) and male (f) mice, showing that all mouse groups lack allodynia. Data are presented as mean ± SEM. Statistical significance (p-values shown) was determined using a 2-way repeated measures ANOVA with Šídák’s multiple comparisons test between Air mCCI vs EtOH mCCI groups. Female - Treatment × Time interaction, F(3.815, 30.52) = 14.89, p < 0.0001; Time, F(3.815, 30.52) = 13.82, p < 0.0001; Treatment F(1, 8) = 178.8, p < 0.0001; Geisser–Greenhouse correction applied ε = 0.4768. Male - Treatment × Time interaction, F(4.537, 58.98) = 21.98, p < 0.0001; Time, F(4.537, 58.98) = 21.06, p < 0.0001; Treatment F(1, 13) = 301.2, p < 0.0001; Geisser–Greenhouse correction applied ε = 0.5671.

We first measured baseline (BL) mechanical sensitivity in both hind paws prior to injury and found comparable thresholds between the EtOH- and air-exposed groups, regardless of sex (**Fig. 6c–f**), indicating equivalent sensory function before surgery. Mechanical withdrawal thresholds were then measured longitudinally every other day for 16 days in both the ipsilateral (injured) and contralateral (uninjured) paw. Following mCCI, both EtOH-exposed males and females rapidly developed robust mechanical allodynia in the injured (ipsilateral) paw by the third day post-surgery, followed by progressive reductions in withdrawal thresholds that persisted throughout the 16-day testing period (Female - Treatment × Time interaction: F(3.815, 30.52) = 14.89, p<0.0001; Male - Treatment × Time interaction: F(4.537, 58.98) = 21.98, p<0.0001) (**Fig. 6c,d**). Consistent with the observed sex-specific behavioral effects of TTAE, EtOH-exposed males developed allodynia more rapidly and exhibited lower response thresholds than EtOH-exposed females, indicating heightened mechanical sensitivity. In contrast, allodynia was not observed in any sham-operated or Air-exposed mCCI groups (**Fig. 6c,d**), nor in the contralateral hind paw of any treatment group (**Fig. 6e,f**).

Overall, these findings demonstrate for the first time that alcohol exposure restricted to the third trimester-equivalent developmental period is sufficient to increase susceptibility to neuropathic pain following peripheral nerve injury. Importantly, the magnitude and persistence of the allodynia response observed in TTAE animals suggest that late developmental exposure may produce sensory vulnerability comparable to, or potentially greater than, that reported in prenatal alcohol exposure models targeting earlier gestational stages.

## 4. DISCUSSION

The present study demonstrates that alcohol exposure restricted to the third trimester-equivalent developmental period is sufficient to produce persistent alterations in behavioral regulation and sensory processing in adulthood. Using a neonatal vapor exposure paradigm that achieved binge-level blood ethanol concentrations, we found that TTAE produced long-lasting changes in locomotor activity, exploratory behavior, recognition memory, risk-related decision making, and susceptibility to neuropathic pain in a sex-dependent manner. These findings extend previous work showing that developmental alcohol exposure produces enduring neurobehavioral consequences and further support the concept that the timing of exposure is a critical determinant of functional outcome, reflecting the stage-specific sensitivity of the developing brain to environmental insults (Guerri et al., 2009; Patten et al., 2014; Noor & Milligan, 2018).

One of the most striking observations in the present study is the absence of a significant effect of TTAE on female locomotor activity despite robust increases in activity observed in males. Baseline locomotor activity levels were inherently higher in females than in males (Lightfoot et al., 2008), suggesting that female locomotor behavior may operate near a physiological ceiling under standard conditions. Under this framework, females may possess limited dynamic range for further increases in activity following developmental perturbation, whereas males begin from a lower baseline and therefore retain greater capacity for alteration of locomotor activity. Sex-dependent behavioral responsiveness to developmental alcohol exposure has been reported in prior studies, with PAE producing differential effects on exploratory and stress-related behaviors in males and females depending on the behavioral context and developmental timing (Osborn et al., 1998). Importantly, the observation that activity levels in TTAE males approached those observed in control females further supports the interpretation that TTAE shifts male behavior toward a higher activity set point rather than producing pathological hyperactivity.

Despite the increase in locomotor activity observed in males, TTAE did not produce overt baseline anxiety-like behavior in the open field task, indicating that the behavioral effects of TTAE reflect altered locomotor regulation and exploratory drive rather than generalized alteration in anxiety. This distinction is important because many earlier studies of PAE as well as TTAE have reported increased anxiety-like behavior characterized by reduced exploration of open or novel environments (Baculis et al., 2015; Jessup et al., 2026; Osborn et al., 1998). These differences suggest that behavioral outcomes following developmental alcohol exposure depend strongly on the stage, duration, and degree of exposure. Early gestational exposure primarily disrupts neurogenesis and neuronal migration, whereas late gestational exposure more strongly affects synaptic maturation, myelination, and circuit refinement, leading to distinct behavioral phenotypes (Guerri et al., 2009; Newville et al., 2017, 2022).

Additionally, the present study revealed selective impairments in novelty recognition in both TTAE and control mice. However, TTAE males demonstrated a significant increased preference for the familiar object over the novel object relative to control males, indicating deficits in recognition memory or novelty processing. Surprisingly, neither the female groups nor the air-exposed control males showed recognition of the novel object based on discrimination index. Although the memory retention deficits or inability to recognize the novel object may be due in part to the 24-hour period between the training and testing sessions, a previous study of neonatal pups intubated with binge-level of ethanol twice daily from P3 to P20 show that the alcohol exposed mice were able to discriminate the novel object from the familiar, albeit at significantly reduced discrimination index (Dursun et al., 2025). These learning differences suggest that the route of alcohol exposure combined with baseline behavioral differences may likely influence behavioral responsiveness in mice.

The elevated zero maze results further illustrate the complex and sex-specific nature of behavioral regulation following TTAE. Despite similar frequency of entries into the open arms of the maze, TTAE males exhibited increased time spent in the open arms, indicative of enhanced risk-taking behavior or reduced behavioral inhibition, whereas females demonstrated the opposite pattern, spending less time in the open arms and exhibiting increased avoidance behavior. This divergence suggests that TTAE does not uniformly increase or decrease anxiety-like behavior but instead alters the balance between exploration and risk evaluation in a sex-dependent manner. Interestingly, earlier studies of both PAE and TTAE mice have frequently reported reduced open-arm exploration and increased anxiety-like behavior for both males and females in the elevated plus maze, indicating a behavioral phenotype somewhat distinct from that observed in the present study (Baculis et al., 2015; Osborn et al., 1998; Patten et al., 2014). However, it is possible that these differences may reflect variation in developmental timing as well as duration and dose of ethanol exposure, which were much lower in BECs in these earlier studies compared to the current study.

An additional and particularly important finding of the present study is the demonstration that TTAE increases susceptibility to neuropathic pain following minor peripheral nerve injury in adulthood. Both male and female mice exposed to alcohol during the third trimester-equivalent period developed robust mechanical allodynia following mCCI, with males exhibiting more rapid onset and greater sensitivity than females. These findings extend previous work demonstrating that PAE primes neuroimmune pathways and increases vulnerability to chronic pain later in life (Noor et al., 2017; Noor et al., 2019). In these studies, PAE was shown to enhance spinal glial activation and inflammatory cytokine signaling following nerve injury, suggesting that early-life alcohol exposure can produce long-term changes in immune and neural signaling pathways that increase susceptibility to pathological pain states (Noor et al., 2017). At the mechanistic level, activation of microglia and astrocytes following ethanol exposure has been identified as a key mechanism underlying persistent changes in neural connectivity and behavioral function (Noor et al., 2018). The neuroimmune proinflammatory profiles and the extent to which microglia and astrocytes are activated in the TTAE mice will need to be investigated in the future.

Taken together, the findings of the present study demonstrate that alcohol exposure during the third trimester-equivalent developmental period produces persistent and sex-dependent alterations in behavioral regulation and sensory processing in adulthood. Collectively, these findings provide new insight into how alcohol exposure across our all trimesters contributes to the persistent functional deficits characteristic of FASDs and underscore the importance of considering biological sex in the evaluation of neurobehavioral risk following PAE.

## AUTHOR CONTRIBUTIONS

E.V., C.F.V., and T.Y.V. designed research; E.V., M.S.S., H.C., L.E.P.B., E.J.B., and T.Y.V. performed research; C.J.M. and R.A.M provide behavioral trainings; E.V., M.S.S., E.D.M., and T.Y.V. analyzed data; E.V., C.F.V., and T.Y.V. wrote the paper. All authors provide approve of the manuscript.

## ACKNOWLEDGEMENT

This research was supported by NINDS R01NS121660 and NIAAA P50AA022534 NMARC Pilot 6B and Component 6 grants (T.Y.V.) and the UNM Center for Brain Recovery and Repair (CBRR) NIGMS P20GM109089 grant. The Preclinical Core in the CBRR, were essential toward the completion of this research.

